# Efficient CRISPR–Cas genome editing in brown algae

**DOI:** 10.1101/2025.07.21.665871

**Authors:** Cláudia Martinho, Masakazu Hoshino, Morgane Raphalen, Viktoriia Bukhanets, Anagha Kerur, Kenny Bogaert, Rémy Luthringer, Susana Coelho

## Abstract

Brown algae represent the third most complex lineage to evolve multicellularity, independently from plants and animals. However, functional studies of their development, evolution, and biology have been constrained by the lack of efficient and scalable genome editing tools. Here, we report a robust, high-efficiency, and transgene-free CRISPR–Cas12-based genome editing method applicable across four ecologically and biotechnologically important brown algal species. Using *Ectocarpus* as a model, we optimized a PEG-mediated RNP delivery system employing a temperature-tolerant Cas12 variant, achieving reproducible, high-efficiency editing across multiple loci without the need for cloning or specialized equipment. As proof of concept, we precisely recapitulated the hallmark *imm* mutant phenotype by editing the IMMEDIATE UPRIGHT (IMM) locus, a phenotype previously described only from a rare spontaneous mutation. APT/2-FA-based selection further improved specificity with minimal false positives. The protocol was readily transferrable to other species, including kelps long considered recalcitrant to transformation. This platform now makes functional genomics accessible in brown algae, enabling mechanistic dissection of developmental processes, life cycle transitions, and the independent origins of complex multicellularity. Our work enables the broader adoption of brown algae as experimental models and provides a valuable platform for marine biotechnology and evolutionary research.

**Motivation:** Although most of biodiversity on Earth lives in oceans, a significant proportion of its organisms remain largely uncharacterized. Brown algae represent one of such understudied group of marine photosynthetic eukaryotes. Despite their importance as emerging models for developmental evolution and blue biotechnology, functional genomics in brown algae has remained largely inaccessible due to a lack of efficient and scalable genome editing tools. Our aim is to democratize genome editing in brown algae by developing a high-efficiency, transgene-free protocol that works across multiple species, without the need for specialized equipment. This high-efficiency method fully enables the field of functional genomics in an unexplored multicellular lineage.

**Highlights:**

- High-efficiency, low-cost genome editing in brown algae without specialized equipment.
- ·Applicable to non-model species, including those of economic importance.

## Introduction

Brown algae represent a lineage of marine photosynthetic eukaryotes that evolved complex multicellularity independently of plants and animals^1^ through mechanisms that remain largely unexplored^2–5^. Ubiquitous in coastal environments, they play key ecological roles, serve as a natural source of bioactive compounds for industrial applications, and constitute an important food resource^6^. However, the lack of efficient and broadly accessible genome editing protocols has hindered both fundamental and applied research.

Previous efforts to deliver Clustered Regularly Interspaced Short Palindromic Repeats (CRISPR)–Cas9 ribonucleoproteins (RNPs) in brown algae employing biolistic and/or microinjection methods have successfully yielded genomic edits^7,8^. However, these labour-intensive approaches rely on specialized equipment, require extensive technical training, and are limited by relatively low editing efficiencies; for instance, biolistic delivery has been associated with a false-positive rate of approximately 50%^7^.

Here, we present a genome editing method based on Polyethylene glycol (PEG)-mediated RNP-transfection that achieves unprecedented efficiency with negligible false-positive rate across four brown algae: *Ectocarpus* sp. 7 and *Scytosiphon promiscuus*, widely used systems for fundamental research^9–12^, *Laminaria digitata,* a laminariacean kelp of ecological and economic importance, and *Undaria pinnatifida* (wakame), an alariacean kelp and one of the most commercially cultivated brown algae worldwide^6^.

## Results

### PEG-mediated RNP *Ectocarpus* gamete transfection allows highly-efficient genome editing

To develop a simple and highly efficient genome editing method in Ectocarpus sp., we exploited its ability to regenerate haploid partheno-sporophytes (PSPs) from unfertilized gametes via parthenogenesis (**Fig. 1A**). Notably, Ectocarpus gametes remain cell wall-free for approximately two hours following release from gametangia in the absence of fertilization^3^, providing a transient window during which CRISPR–Cas RNPs can be efficiently transfected.

**Figure 1.**
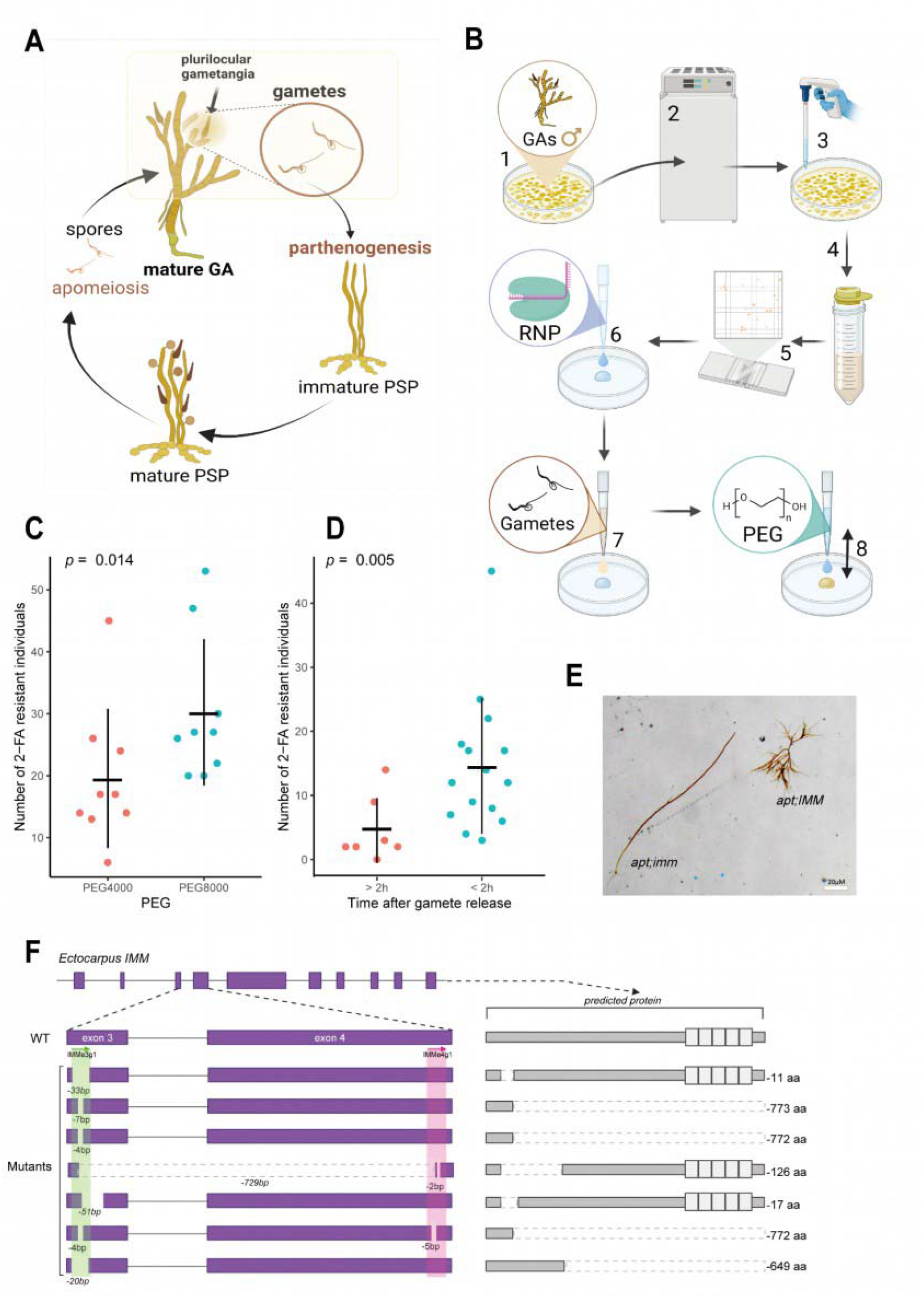
*Ectocarpus* CRISPR-based genome editing. **A)** Parthenogenetic life cycle of *Ectocarpus* sp. Gametophyte (GA) produces gametes which parthenogenetically develop into partheno-sporophyte (PSP). **B)** Schematic diagram of the process of the PEG transfection for *Ectocarpus sp*. 7 The process begins with gametophytes (GAs) cultured in Petri dishes (step 1), followed by incubation to induce gamete release (step 2). The gametophytes are then transferred using a pipette (step 3), filtered to separate from gametophytic tissue, and centrifuged to concentrate gametes (step 4). After centrifugation, the concentration of gametes is examined using a hemocytometer (step 5). Ribonucleoprotein (RNP) is added to the petri dish (step 6) and mixed with gamete suspension (step 7). Finally, 40% w/v polyethylene glycol (PEG), prepared in seawater, is applied to facilitate transfection and vigorously mixed by pipetting with a wide bore tip (step 8). See details in methods section. **C)** Comparison of the number of 2-FA resistant individuals following treatment with different PEG molecular weights: PEG4000 and PEG8000. Dots depict independent trials (Table S2). The horizontal and vertical lines in the scatter plot represent the mean and standard deviation, respectively. Significantly more resistant individuals were observed with PEG8000 treatment (Exact Wilcoxon rank sum test, *p* = 0.014). **D)** Comparison between PEG treatments using gametes collected more than 2 hours after release (>2h) and those using freshly released gametes (<2h). A significantly higher number of resistant individuals was observed when fresh gametes were used (exact Wilcoxon rank sum test, *p* = 0.005). **E)** Morphological difference of double KO mutant of *APT* and *IMM* (*apt;imm*) and *APT* single KO mutant (*apt;IMM*). The *APT* single KO mutant develops prostrate filaments, whereas the double KO mutant develops long upright filaments. **F)** Schematic representations of the *Ectocarpus IMM* gene model of WT and mutant individuals. Exon regions are shown as purple boxes, and crRNAs are indicated with arrows. The expected protein products for the WT and each mutant are shown to the right of their respective gene models. The five imperfect tandem repeats of a 38 amino acid cysteine-rich motif, characteristic of the IMM C-terminal region, are represented as white boxes.

To transfect Cas12-RNPs we employed PEG-mediated RNP transfection, previously successful in the green alga *Ulva prolifera* ^14^. While mature male Ectocarpus gametophytes (GAs) release abundant swimming gamete^15,16^ (**Fig. 1 A**) and were therefore used to optimize the protocol, mutations can be easily backcrossed into wild-type females^15,16^ provided fertility is not affected. Because the optimal growth temperature of this alga is 14°C, we used an engineered Cas12 (Alt-R L.b. Cas12a [Cpf1], IDT) version with increased temperature tolerance and optimal for systems requiring lower culture temperatures. We isolated male gametes from cultivated algae and transfected them with Cas12-RNPs using 40% w/v PEG (Fig. 1B, see methods). As selection marker, we used Cas12 loaded with a APT locus crRNA (Ec-28_000520, Table S1) to generate apt loss-of-function mutations which enable survival in selective medium supplemented with 2-fluoroadenine (2-FA)^7^.

Initially, we tested the effect of an overnight heat-shock at 22°C in darkness after PEG transfection (**Fig. S1A**) and of different Cas12 to crRNA ratios (**Fig. S1B**) with no statistically significant effect on the number of 2-FA resistant obtained (**Fig. S1, Table S2**). However, the heat-shock step was maintained since it resulted in higher number of 2-FA– resistant individuals (**Fig. S1A**). Additionally, we tested the effect of different PEG molecular weights and observed that PEG8000 resulted in higher number of 2-FA resistant individuals (0.003% of transfection efficiency) averaging 30.3 2-FA resistant individuals per trial *i.e.* per Petri dish (**Fig. 1C, Table S2**). Data analysis from multiple independent experiments during protocol optimization, revealed that transfection within the first two hours after gamete release significantly improves editing efficiency (**Fig. 1D, Table S2**), and thus it is recommended to proceed with transfection as soon as gamete release is completed.

Given the method’s high efficiency in generating 2-FA resistant individuals, we next tested whether it could produce double mutants targeting both APT and a gene of interest. To achieve this, we designed two Cas12 crRNAs targeting exons 3 and 4 of the *IMMEDIATE UPRIGHT* (*IMM, Ec-27_002610.1*) gene and co-transfected them with the *APT* RNP complex (**Table S1**). Mutations in the *IMM* locus are known to disrupt basal cell development and accelerate upright filament formation in *Ectocarpus* sp.^5^ providing a clear developmental phenotype to assess the efficiency of our method (Fig. 1E). Remarkably, we observed this characteristic *imm* phenotype in 19 individuals out of 59 (32%) of 2-FA resistant individuals following co-transfection. We selected 15 individuals displaying the *imm* phenotype and confirmed, employing Sanger sequencing, that all harbored mutations at both the *APT* and *IMM* loci (**Table S3**). In contrast, three 2-FA resistant individuals with a wild-type phenotype showed mutations only at the *APT* locus (**Table S3**). Notably, one of the 15 double mutants carried a 723 bp deletion between the two *IMM* target sites, while others had smaller indels at individual sites (**Fig. 1F, Table S3**). While crRNA targeting exon 3 produced mutations in all tested individuals, crRNA targeting exon 4 generated mutations in only 2, suggesting that crRNA or target site susceptibility may differ. Importantly, no false positives were observed since all algae that grew on selective medium and tested, carried mutations in *APT*, confirming the high precision and efficiency of the co-transfection protocol.

### PEG-mediated RNP transfection results in high-frequency genome edits across multiple brown algal species

To test the transferability of the PEG-mediated RNP transfection protocol beyond *Ectocarpus*, we applied the method to other non-model brown algae. Specifically, we transfected *APT*-targeting RNPs into *S. promiscuus* and the kelps *L. digitata* and *U. pinnatifida* (**Fig. 2**), introducing only minor protocol modifications (see Methods). In *S. promiscuus*, as in *Ectocarpus*, gametes released from laboratory-cultivated algae were used for PEG transfection (**Fig. 2A, B**). For the kelps, we instead used meiospores transformation (**Fig. 2C, D**), since kelp gametes exhibit limited parthenogenetic capacity. Meiospores develop into haploid gametophytes, allowing direct phenotypic evaluation of mutation effects without the need for backcrossing or generating homozygous lines.

**Figure 2.**
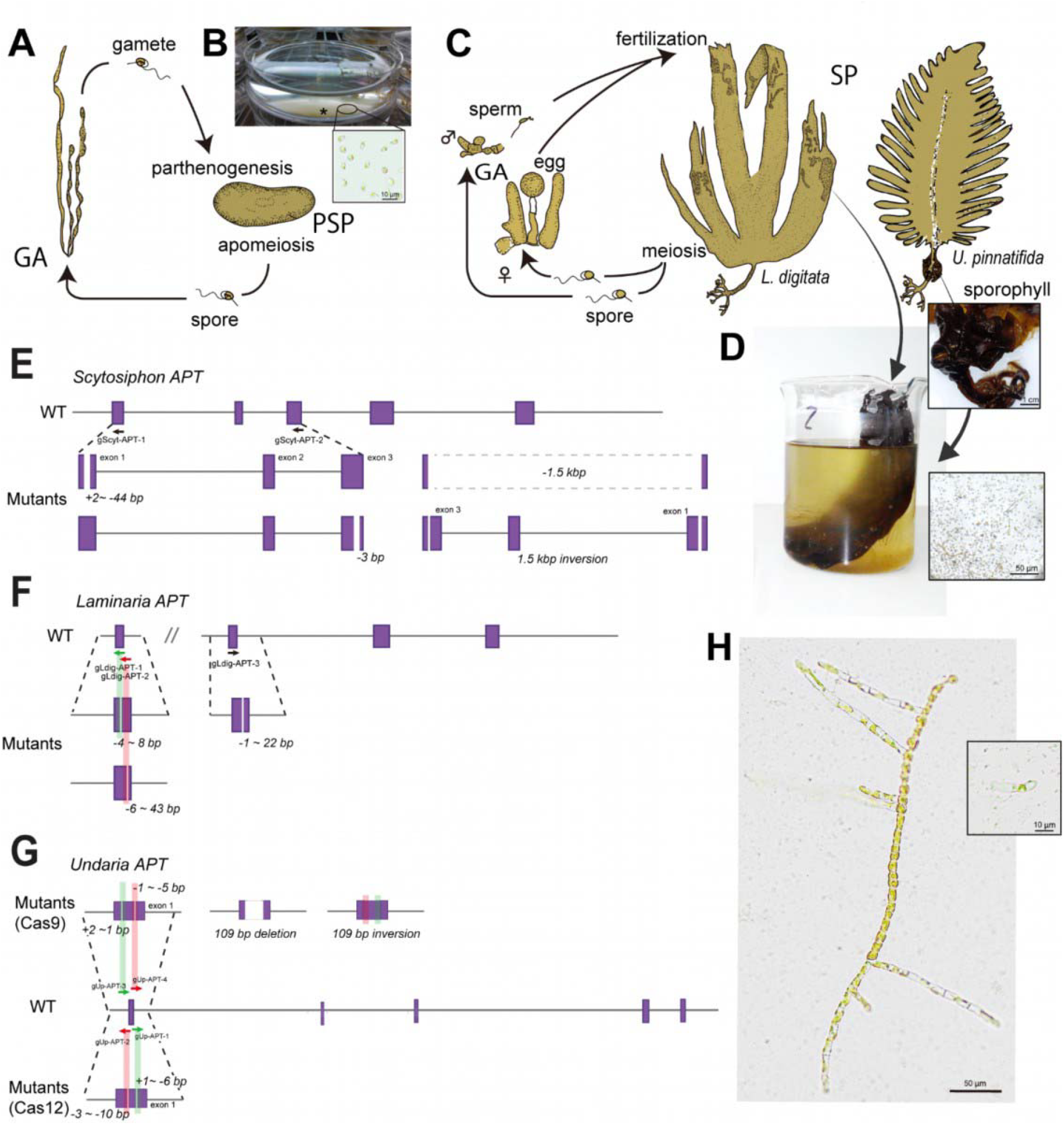
*Scytosiphon* and kelp CRISPR-based genome editing. **A)** Parthenogenetic life cycle of *Scytosiphon promiscuus*. Macroscopic gametophytes (GA) alternate with microscopic discoidal parthenosporophyte (PSP). **B)** Suspension of male gametes of *S. promiscuus* showing accumulation of gametes (asterisk) by negative phototaxis. **C**) Life cycle of *Laminaria digitata* and *Undaria pinnatifida*. In both species, microscopic gametophytes (GA) alternate with macroscopic sporophyte (SP: left, *Laminaria* sporophyte; right, *Undaria* sporophyte). **D)** Released meiospores from *Laminaria* sori and *Undaria* sporophylls. **E)** Schematic representations of the *S. promiscuus APT* gene model of WT and mutant individuals. See details in **Fig. 1. F)** Schematic representations of the *L. digitata APT* gene model of WT and mutant individuals. **G)** Schematic representations of the *U. pinnatifida APT* gene model of WT and mutant individuals following Cas9 transformation (above) and Cas12 transformation (below). **H)** One-month-old germlings of *S. promiscuus APT* mutant and WT (right top) in selective medium.

The transfection rate in *S. promiscuus* was approximately 0.25% allowing to retrieve between 107 and 2458 2-FA resistant individuals with no false positives in the all individuals tested (**Table S4).** In *L. digitata* transformation efficiency varied between 0-0.79% and yielded between 0 and 6128 2-FA resistant individuals, depending on the trial (**Table S4**). This discrepancy was likely due to the fact that fresh zoospores - i.e., actively swimming, wall-less cells-were not available in trials 1 and 3, and instead non-motile, possibly cell wall-bearing spores were used (**Table S4**). In *U. pinnatifida* the transfection efficiency varied among trials: 0.014-0.021%, which reflects more than 100 2-FA resistant individuals per trials. In *L. digitata*, genotyped 2FA-resistants exhibited mutations in the *APT* locus with a false positive rate of 9.3±17.4% among the trials (mean±SD; **Table S4**), while in *S. promiscuus* and *U. pinnatifida*, no false positives could be identified.

For *S. promiscuus*, two *APT*-crRNAs were used simultaneously, causing insertions or deletions (**Fig. 2E**). Most mutants had only one mutation near the crRNA target sites, whereas in two cases, approximately 1.5 kb between the two target sites was entirely deleted (**Fig. 2E**). Additionally, in one case, an inversion occurred within the region between the two target sites (**Fig. 2E**). In *L. digitata*, where only a single crRNA was used for each PEG transfection, only deletions (−1 to −43 bp) were observed (**Fig. 2F**). For *U. pinnatifida* (growing at 20°C) both Cas9 and Cas12 were assayed with each two crRNAs. All were shown to be functional (**Fig. 2G**) with a higher mutation efficiency assessed for Cas12 (**Table S4**). Overall, our results support that PEG-mediated Cas12-RNP complex transfection in gametes or meiospores can be easily scalable and applicable to multiple brown algae species, including kelps of high economic and ecological importance.

## Discussion

Our method enabled efficient genome editing in four brown algal species using either gametes or meiospores. Since most brown algae produce parthenogenetic gametes, and even species without such gametes typically generate meiospores, our approach can be applicable to most brown algal species, provided they produce sufficient numbers of wall-less zoids. In previous studies using biolistic delivery and microinjection in the model algae *Ectocarpus* sp 7, only an average of 0.4 and 1 *apt* mutants were obtained per experiment, respectively^7^, whereas our method can generate on average 30 individuals per trial. In some instances, even higher numbers have been isolated, which we speculate to be related to the quality and viability of the gametes isolated in each trial. Moreover, with biolistic transformation, fewer than half of the 2-FA resistance individuals carried mutations in the *APT* gene, indicating a substantial proportion of false positives^7^ which would require more intensive screening efforts than our method. In contrast, our method produced tens to thousands of 2-FA resistance individuals per experiment across four different brown algal species, with false positives being absent in most experiments and remaining below 10% at highest. By enabling precise, efficient, and accessible genetic manipulation, our approach opens the possibility to explore the molecular mechanisms underlying brown algal development, physiology, and evolution. Its high efficiency combined with high reproducibility and easy implementation, will enable brown algae studies across a wide community of researchers. The improvement of the protocol therefore represents a major step forward for both fundamental research and biotechnological applications in this unique lineage of complex multicellular marine organisms.

## Limitations of the study

The effectiveness of our method may be limited in Fucales and Dictyotales since these groups lack gametes with parthenogenetic capacity^17^. Additionally, Fucales do not produce meiospores and thus is not possible to circumvent the lack of parthenogenic gametes while Dictyotales produce meiospores that are large and rich in cytoplasm^17,18^ which could potentially interfere with transfection efficiency. These limitations could be circumvented by using brown algae protoplasts for PEG-mediated transfection, as originally reported in tobacco plants^19^ with the caveat that protoplast regeneration is time-consuming^20,21^. Another possibility would be to transform male gametes and perform a cross immediately after transfection, which would then require further generations to isolate homozygous individuals. While microinjection is labor-intensive and requires specialized equipment, introducing RNPs into vegetative cells or embryos in Fucales and Dictyotales remains an alternative for these brown algal groups.

## Acknowledgments

We thank Dr. Kensuke Ichihara for valuable advice. We thank Agnes Henschen for help with DNA extractions, and Andrea Belkacemi and Dorothe Koch for assistance with algal cultures. This study was funded by the Max Planck Society, the European Research Council grant 864038 (SMC), the Bettencourt Foundation (SMC) and the Moore Foundation (SMC). M.H. was supported by JSPS KAKENHI Grant number 23K19386, JSPS Overseas Research Fellowships, and Max Planck Partner Groups.

## Author contributions

MH, CM, SMC conceived and designed the experiments. CM, VB, MH and MR developed protocols. CM, VB, MH, AK, KB, RL and MR performed experiments. MH, CM, SMC wrote the manuscript with help of all authors. All authors provided critical feedback and helped shape the research, analysis and manuscript writing.

## STAR Methods

### *Ectocarpus* sp.7 PEG-mediated RNP transfection

*Ectocarpus* sp.7 gene sequences were retrieved from v3 reference genome^22^ and CRISPOR^23^ used to design Cas12 guide RNAS with limited off-targets and highest efficiency scores. Male gametophytes (strain Ec32) were cultured in plastic Petri dishes (150 × 15 mm) (10 individuals per Petri dish) and sterile natural sea water (NSW) enriched with half-strength Provasoli medium^24^ (Provasoli enriched sea water: PES) at 14°C in neutral day (12 h: 12 h, light:dark) conditions with LED lighting of 20 μmol m^−2^ s^−1^ photon flux density. Male gametophytes were isolated from individual unilocular sporangia, which in turn were isolated from Ec32 mature partheno-sporophytes (3-4-week old), as previously described^25^. Mature gametophytes (displaying clear accumulation of mature plurilocular sporangia) are observed usually after 3-4 weeks after culture preparation and clearly visible under a light stereoscope^4,25^. Gametophytes grown on 50 Petri dishes (500 gametophytes) were collected under a laminar flow hood on a sieve with a 50 μm mesh and rinsed with NSW at 14°C to remove small filaments and already released gametes. The biomass was concentrated and equally distributed in small Petri dishes (60×15mm) up to 100 gametophytes per Petri dish and the excess of water was removed with a 200 μL pipet. To maintain high humidity levels a few (typically 4-8) drops of NSW were added on the edge of the Petri dish. The dishes were sealed with parafilm, wrapped in aluminium foil to keep darkness conditions and transferred for 3 hours to 14°C. Gamete release was induced by adding 5 mL of 4°C NSW and incubating 5 minutes at room temperature under 40 μmol m^−2^ s^−1^ photon flux density. Gametophytes were incubated further 30 minutes at 14°C with LED lighting of 40 μmol m^−2^ s^−1^ photon flux density to allow for further gamete release. The gametes were then separated from gametophytic tissue by filtering through a 10 μM cell strainer into a 50 mL conical plastic tube and concentrated by centrifuging in a swing-out rotor (4600 rpm) for 5 minutes at Room Temperature (RT). The majority of the supernatant was removed by pipetting and 200-300 μL of NSW were left to resuspend the gametes. The gamete suspension was then diluted 1:10 and stained with 1 μL of lugol solution to count in a hemocytometer. The gametes were then diluted to 1×10^4^/μL in NSW. For PEG-mediated transfection, in a laminar flow hood, 20 μL of RNP complex mixture (or 20 μL mock control with IDT buffer) were pipetted into a plastic Petri dish (90×20mm) and 100 μL of the gamete suspension (10^6^ gametes) were gently mixed by pipetting. This gamete/RNP solution was then mixed with 120 μL of 40% w/v PEG-8000 (Fisher Scientific, Schwerte, Germany) or PEG-4000 (Sigma, Darmstadt, Germany) solution, filtered through a 0.22 μM filter by pipetting up and down. PEG pipetting and mixing was carried out vigorously with a wide bore 200 μL filter tip to ensure proper homogenization. A small degree of bubbling is expected and did not, in our hands, influence transformation efficiency. The gametes were then incubated at RT in darkness for 30 minutes. The 20 μL of the RNP complex was prepared as following: 1.2 μL of guide RNA (100 μM, IDT, Leuven, Belgium), 2.8 μL of IDTE Buffer (IDT), 8 μL of Alt-R L.b. Cas12a (Cpf1, 15.6 μM, IDT), 8 μL of 2.5X NEB Buffer 3.1 (New England Biolabs, Ipswich, MA, USA). For transfections involving multiple RNP complexes, each complex was prepared separately, then combined in the Petri dish, the amount of PEG solution was then adjusted accordingly to maintain a 1:1 ratio (Gamete/RNP:PEG). Following transfection, 20 mL of half-strength PES was added to each plate to stop transfection and enable gamete germination. Dishes were incubated overnight in the dark at 22°C (typically 16 hours), then transferred the next morning to 14°C in neutral day conditions (12 h: 12 h, light:dark). 48h after transformation, 2-fluoroadenine (2-FA, Sigma) was added to the Petri dishes to a final concentration of 20 μM, and samples were incubated in a in neutral day conditions (12 h: 12 h, light:dark) with LED lighting of 20 μmol m^−2^ s^−1^ photon flux density until the isolation of 2-FA resistant algae and therefore bearing putative mutations in *APT* and the gene of interest.

Six to eight weeks after the transfection, the number of growing germlings in 2-FA supplemented culture medium (putative *apt* mutants) were counted, and some of them were carefully isolated to 2-FA free half-strength PES using forceps to generate enough biomass for genotyping and validation Genotyping PCR was performed using the Terra PCR Direct Polymerase Mix (Takara) or the Kapa G3 Plant PCR kit (RocheBiosystems) with 2uL of the following suspension: small fragments of algal tissue (approximately 1mm^2^) grinded in 60μL of Nuclease free Water (Ambion). Sanger sequencing was performed by Azenta. Details of gRNA and primers are provided in **Table S1.**

### *Scytosiphon promiscuus* PEG transfection

The *APT* gene of *S. promiscuus* was identified by aligning DNA sequencings of the *Ectocarpus sp.7 APT* gene against the *S. promiscuus* genome^26^ using DIAMOND2^27^. A single gene (gene code: mRNA_S-promiscuus_M_contig7.17323.1) on chromosome 28 was identified as *APT* gene of *S. promiscuus*.

Male gametophytes (strain As6m) were cultivated in plastic Petri dishes (90 × 20 mm) and sterile NSW enriched with full strength PESI medium^28^ at 10°C in long day (16h:8h, light:dark) conditions with LED lighting of 20 μmol m^-2^s^-1^ photon flux density. Medium was renewed every week until gamete collection. Mature gametophytes release gametes immediately after the medium renewal and thus cultures were closely inspected after each media change. Because *S. promiscuus* gametes have negative phototaxis, freshly released gametes accumulate on the opposite side of a light source in a Petri dish. The accumulated gametes were collected and diluted to 0.4–10 million per 1 mL with NSW. For PEG transfection, 100 μL of the gamete suspension was gently mixed with 8 μL of RNP complex and 108 μL of 40% w/v PEG-4000 solution in a plastic Petri dish (90 × 20 mm), and then, incubated at room temperature in dark for 30 minutes. To prepare 8 μL of the RNP complex, two different guide RNAs were mixed as following:1 μ□ of guide RNA1 (100 μM), 1 μ□ of guide RNA2 (100 μM), 1.5 μ□ of Alt-R L.b. Cas12a (Cpf1, 67 μM), 3.2 μ□ of 2.5X NEB Buffer 3.1, and 1.3 μ□ of water. After the transfection, 50 mL of full-strength PESI was added to the Petri dishes and incubated in a 22°C dark condition for 48 hours. Then, 2-FA was added to the Petri dishes (final 2-FA concentration of 20 mM), and the samples were incubated in a 14°CC neutral day condition until the isolation of putative *APT* mutants. One month after the transfection, the number of growing germlings in 2-FA supplemented culture medium (putative *apt* mutants) were counted, and eight of them were isolated using glass Pasteur pipets, before being genotyped to validate the mutations. Genotyping was performed as in *Ectocarpus*. Details of gRNA and primers are provided in **Table S1**.

### *Laminaria digitata* PEG transfection

The *APT* gene of *Laminaria. digitata* was identified by aligning *Ectocarpus* sp. 7 *APT* gene against *L. digitata* genome^26^, as described above. The *APT* gene of *Laminaria* (gene codes: mRNA_L-digitata_M_contig27133.1.1 and mRNA_L-digitata_M_contig6833.2.1) was divided into two contigs (contig27133 and contig6833) because of the fragmented reference genome assembly.

Sporophytes of *L. digitata* were collected at Santec, France on 8 December 2022. The sporophytes were washed with sterile NSW and pat dried to remove water. The sori were excised from the sporophytes and transferred to the laboratory in a chilled transport box. To induce meiospore release, the sori were placed in fresh sterile NSW at room temperature one day after the sampling. However, this did not result in immediate meiospore release. The sori were therefore incubated in sterile NSW in 14°C neutral day conditions until meiospores were released up to two days. Meiospores were collected from three independent sporophyte individuals. Meiospore release resulted in a change in water colour towards brown tones. This brown seawater was transferred to 50 mL tubes and centrifuged at 4000 g for 1 minute. After the centrifugation, the supernatant was removed and fresh sterile seawater was added to the pellet of meiospores and the density of meiospores was adjusted to 0.54–7.7 million per 1 mL. PEG transfection was performed as in *S. promiscuus*, except that each guide RNA was used independently. The RNP complex were prepared as following: 1 μL of guide RNA (100 μM), 0.75 μL of Alt-R L.b. Cas12a (Cpf1, 67 μM), 3.2 μL of 2.5X NEB Buffer 3.1, and 3.05 μL of water. After the transfection, 50 mL of full-strength PESI was added to the Petri dishes and incubated in a 22°C dark condition for 48 hours. Then, 2-FA was added to the Petri dishes to reach a concentration of 20 mM and incubated in a 14°C neutral day condition (green light) until the isolation of putative *apt* mutants. Some of these putative mutants were isolated and genotyped to confirm the mutations. Genotyping was performed as in *Ectocarpus*. Details of gRNA and primers are provided in **Table S1.**

### *Undaria pinnatifida* PEG transfection

The *APT* gene of *Undaria pinnatifida* was identified by aligning *Ectocarpus* sp. 7 *APT* gene against the *U. pinnatifida* genome^29^, as described above. The *APT* gene of *Undaria* (UNPIN0032CG0760) is located on contig LG22.

Mature sporophytes of *U. pinnatifida* were obtained by crossing the male gametophyte strains Un1f and Un1m, isolated from a single sporophyte (Bizeux, St. Malo, France) cultured in sterile NSW enriched with half-strength PES under controlled laboratory conditions (16–20°C, 16:8 h light:dark) with LED lighting at 10 μmol□m□^2^□s□^1^ photon flux density. Spore release was induced by transferring sporophyll tissue to fresh NSW and incubating at room temperature with gentle shaking for 20 minutes. The spore suspension was then collected and its concentration was measured using a haemocytometer. A suspension of approximately 7.5×10□ spores per 100 μL was used for transfection.

For PEG transfection, 100 μL of the spore suspension was gently mixed with 20 μ□ of RNP complex and 120 μ□ of 40% w/v PEG-8000 (Fisher Scientific, Schwerte, Germany), filtered through a 0.22 μm syringe filter, in a sterile plastic Petri dish (90 × 20 mm), and then incubated at room temperature in the dark for 30 minutes. 20□μ□ of RNP complex was prepared as follows: for Cas9, 4 μ□ of crRNA:tracrRNA duplex (100 μM each, IDT, Leuven, Belgium), annealed in IDT Duplex Buffer, 8 μL of Cas9 enzyme (15.6 μM, IDT), and 8 μL of 2.5X NEB 3.1 Buffer (New England Biolabs, Ipswich, MA, USA); for Cas12a, 4 μ□ of crRNA (100 μM, IDT Leuven, Belgium), 8 μ□ Alt-R L.b. Cas12a (Cpf1,IDT), and 8 μL of 2.5X NEB 3.1 Buffer. For dual RNP transfections, each complex was prepared separately and combined in the dish prior to PEG addition.

Following transfection, 20 mL of sterile NSW supplemented with half-strength PES was added to each dish. The plates were incubated overnight at room temperature in the dark and then transferred the next morning to normal culture conditions at 20°C with a 16:8 light:dark cycle.

Forty-eight hours after transformation, 2-fluoroadenine (2-FA, Sigma) was added to each plate at a final concentration of 75□μM. Samples were incubated under standard growth conditions with LED lighting of 10–20 μmolLmL^2^LsL^1^ photon flux density for one week. Surviving 2-FA-resistant gametophytes were isolated and grown individually for further genotyping to validate *APT*-targeted mutations using Sanger sequencing as in *Ectocarpus*. Details of gRNA and *APT* primers are provided in **Table S1**.

## Supplementary Figures

**Figure S1.**
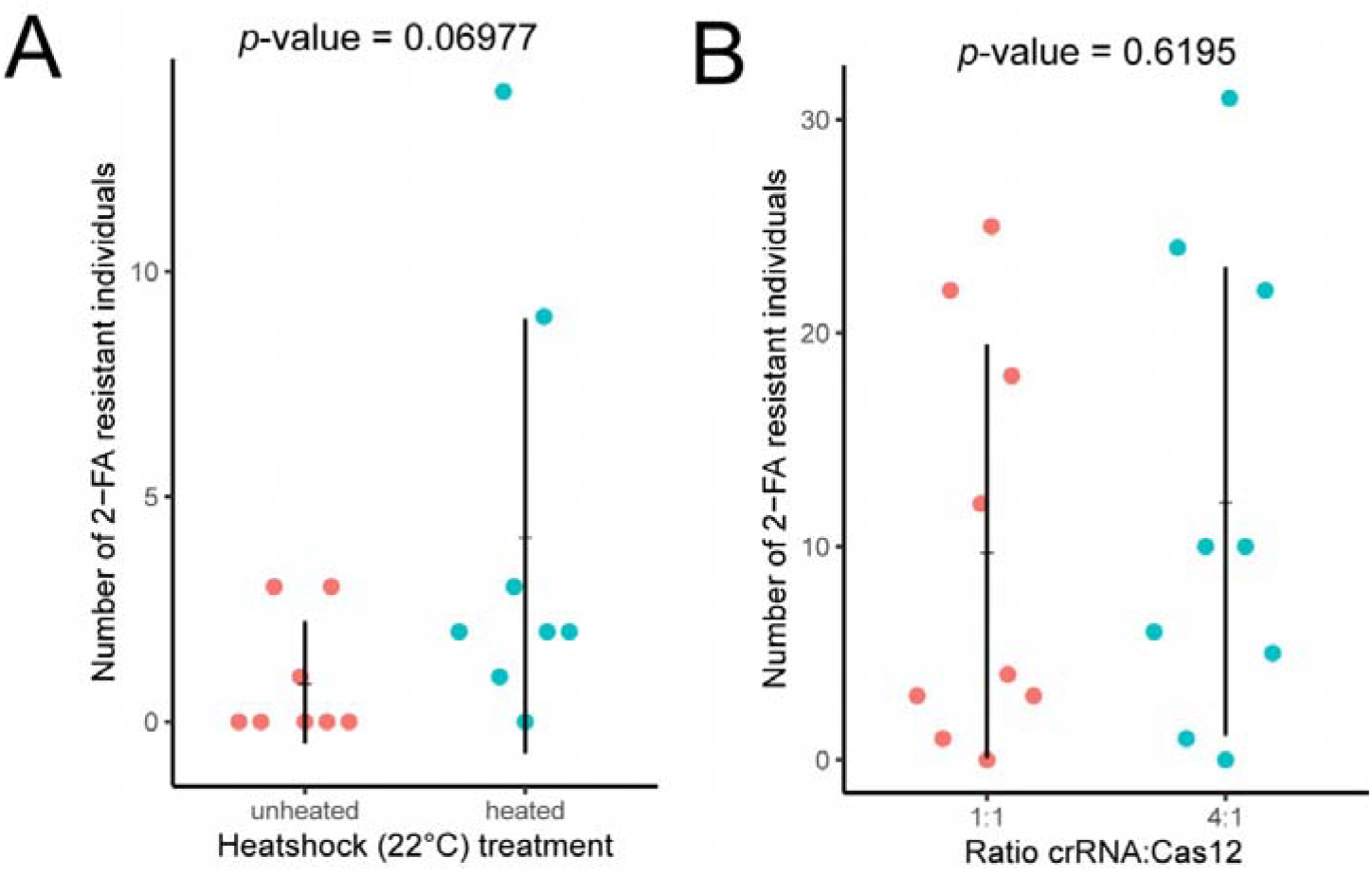
Protocol optimization for *Ectocarpus*. **A)** Scatter plot showing the effect of the heat shock at 22ºC on the number of 2-FA resistant individuals. The horizontal and vertical lines in the scatter plot represent the mean and standard deviation, respectively. No significant effect was detected (exact Wilcoxon rank sum test, *p* = 0.06977). **B)** Scatter plot showing the effect of the crRNA:Cas12 ratio (1:1 vs. 4:1) on the number of 2-FA resistant individuals. No significant effect was detected (exact Wilcoxon rank sum test, *p* = 0.6195).

## Table Legends

Table S1. crRNAs (guide RNAs) and primers used in the present study.

Table S2. Result summary of the optimization of PEG condition for *Ectocarpus*.

Table S3. Results summary of *IMM* mutagenesis for *Ectocarpus*.

Table S4. Result summary of the PEG transfection for *Scytosiphon, L. digitata*, and *U. pinnatifida*.

